# Octopamine integrates the status of internal energy supply into the formation of food-related memories

**DOI:** 10.1101/2023.06.01.543187

**Authors:** Michael Berger, Michèle Fraatz, Katrin Auweiler, Katharina Dorn, Tanna El Khadrawe, Henrike Scholz

## Abstract

The brain regulates food intake in response to internal energy demands and food availability. However, can internal energy storage influence the type of memory that is formed? We show that the duration of starvation determines whether *Drosophila melanogaster* forms appetitive short-term or longer-lasting intermediate memories. The internal glycogen storage in the muscles and adipose tissue influences how intensely sucrose associated information is stored. Insulin-like signaling in octopaminergic reward neurons integrates internal energy storage into memory formation. Octopamine, in turn, suppresses the formation of long-term memory. Octopamine is not required for short-term memory, because octopamine-deficient mutants can form appetitive short-term memory for sucrose and to other nutrients depending on the internal energy status. The reduced positive reinforcing effect of sucrose at high internal glycogen levels combined with the increased stability of food-related memories due to prolonged periods of starvation could lead to increased food intake.

## Introduction

To ensure survival, the internal energy status of an animal needs to be adjusted to the energy expenditure of the organism and the availability of external food. Increased storage of energy correlates with increased food intake in the past. Dysregulation of food intake can lead to diseases such as obesity and diabetes. Sucrose is a carbohydrate enriched in Western diets, and the breakdown product of sucrose - glucose - can be stored in the organism as glycogen, primarily in the liver and muscles. Elevated glycogen levels are a hallmark of glycogen storage diseases that are accompanied by defects in the liver, muscles and brain (Ellingwood and Cheng, 2018). Similar to vertebrates, the fruit fly *Drosophila melanogaster* uses glucose as its primary energy source and stores glycogen primarily in the muscles—the main site of energy expenditure—and the fat body—the equivalent of the vertebrate liver (Galikova and Klepsatel, 2023; Wigglesworth, 1949). As in vertebrates, glycogen levels are also found in the brain (Yamada et al., 2018).

In *Drosophila*, the monoamine octopamine - functionally related to noradrenalin in vertebrates - is involved in the regulation of energy homeostasis. In tyramine-β-hydroxylase (*Tβh*) mutants lacking the neurotransmitter octopamine, the glucose and trehalose concentrations in the hemolymph change less upon starvation, and the life span is extended (Damrau et al., 2017; Li et al., 2016). *Tβh* mutants show an increased threshold to respond to sucrose with the extension of their proboscis when sucrose is offered to their tarsi – a structure that contains gustatory receptor neurons. The reduced responsiveness to sucrose correlates with a reduced sucrose intake in *Tβh* mutants (Li et al., 2016; Scheiner et al., 2014). Octopaminergic neurons potentiate the response of sugar-sensing gustatory receptor neurons in satiated flies, suggesting that octopaminergic neurons regulate feeding behavior by changing the sensitivity of taste receptor neurons (Youn et al., 2018). The reduced responsiveness to sucrose is also thought to be responsible for defects observed in habituation in *Tβh* mutants (Scheiner et al., 2014). In addition to their changes in simple forms of neuronal plasticity, *Tβh* mutants fail to form a positive association with a food reward in a classical olfactory conditioning paradigm. Flies quickly learn to associate an odorant with sucrose, and T*β*h mutants do not show this positive association directly after learning (Schwaerzel et al., 2003). However, when a longer time after training has elapsed, memory appears (Das et al., 2014). The release of octopamine mediates the reinforcing effect of sweet taste in short-term memory and the anesthesia-resistant form of long-term memory (Burke et al., 2012; Huetteroth et al., 2015; Wu et al., 2013). The question arises how the internal energy supply is integrated into the formation of food related memories and feeding behavior in *Tβh* mutants.

The peptide hormone insulin regulates glycose homeostasis at the cellular level (Saltiel and Kahn, 2001). Insulin and its *Drosophila* counterpart, insulin-like peptides, perform their functions through well-conserved signal transduction cascades (Chatterjee and Perrimon, 2021; Inoue et al., 2018). In vertebrates, in addition to its function in fat tissue and muscles, the insulin receptor is broadly expressed in the brain and regulates neuronal plasticity (Nakai et al., 2022). For example, in rats, reduced insulin receptor function in the hypothalamus results in loss of long-term potentiation and impaired spatial memory (Grillo et al., 2015). In *Drosophila*, the insulin receptor substrate Chico is required for the development of mushroom bodies, a brain region required for learning and memory, and loss of Chico results in learning defects (Naganos et al., 2012). Disrupting the function of the insulin receptor in the mushroom bodies also results in learning defects, whereas the insulin receptor and Chico are required in the ellipsoid body for protein biosynthesis-dependent long-term memory (Chambers et al., 2015). While there is evidence that insulin receptor signaling regulates cellular physiology, it is not clear how the internal energy supply of the animal influences the formation of food-related memories.

To analyze the relationship between energy status, the evaluation of a food reward and the formation of memory and food intake, we used the olfactory associative learning and memory paradigm. Using nutrients as a positive reward allows for investigating how the internal energy status influences the evaluation of the reinforcer and in turn learning and memory. As a step to understand the interconnection between reward evaluation, nutrient intake and the formation of food-related memories, we analyzed the function of octopamine in the regulation of internal energy homeostasis, learning and memory and food intake.

## Results

Starvation increases learning and memory performance when using a food-related reward (Colomb et al., 2009). However, how the internal energy status is integrated into the evaluation of the reward and the stability of food-related memories and whether there is a correlation between the perceptions of the food reward, food-related memories and food intake are not clear.

### The duration of hunger influences the strength and stability of memories

To investigate whether changes in internal energy storage influence food-related memories, we used the olfactory associative learning and memory paradigm using sucrose as a food-related reinforcer (Schwaerzel et al., 2003). In the paradigm, flies learn to associate an odorant with sucrose and are later tested on whether they remember the association. A prerequisite for this association to be made is that the animals are hungry. During the training, starved flies were exposed for 2 min to the unconditioned odorant, 4-methylcyclohexanol (MCH), followed by a 2 min exposure to a second odorant, 3-octanol (3-OCT), that was paired with the reward of 2 M sucrose. After the training, flies were given a choice between the two odorants (Figure 1A). Normally, flies prefer the rewarded odorant already 2 min after the training. To avoid a bias towards one odorant, in a reciprocal experiment, the first odorant was rewarded. The performance index presents the average positive association between the reward and both odorants and describes whether information is learnt and remembered. To ensure that observed differences in learning and memory were not due to changes in odorant perception, odorant evaluation or sucrose sensitivity, different fly populations of the same genotypes were tested for their odorant acuity, odorant and sucrose preference (Table S1). The flies of the different genotypes sensed the odorants and evaluated them as similar salient in comparison. This is important to a avoid bias where flies have to choose between the two odorants after training. They also sensed sucrose. We next determined whether differences in sucrose preference influence sucrose intake during training. Therefore, we measured the intake of food-colored sucrose of starved flies in the behavioral set up. After 2 min, we evaluated whether there was dyed food in the abdomen of the fly (Table S1). Flies of both genotypes fed sucrose within 2 min and there was no difference between *w^1118^* and *Tβh^nM18^* flies.

**Figure 1.**
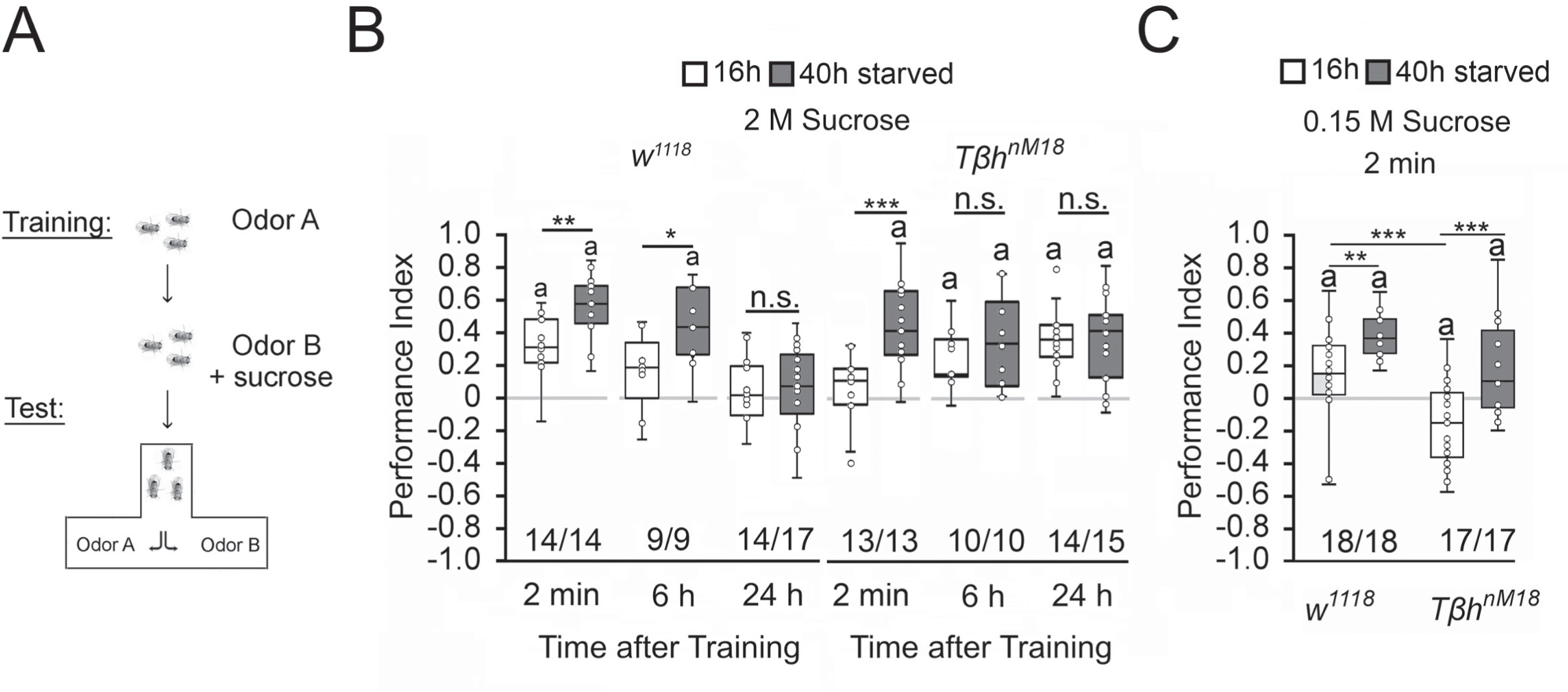
Starvation influences the strength of memory. **A**, Appetitive olfactory learning and memory paradigm. **B**, Appetitive 2 min STM, 6 h and 24 h memory of *w^1118^* or *Tβh^nM18^* that were starved either for 16 h (white bars) or 40 h (dark gray bars) before the training. 2 M sucrose was used as reward. **C**, 0.15 M sucrose was used as reinforcer. Prolonged starvation increases memory performance. Prolonged starvation from 16 h to 40 h leads to the formation of STM in *Tβh^nM18^*. Independent of the duration of starvation, 6 h after training, memory appeared in the mutants. **C,** *Tβh^nM18^* mutants starved for 16 h developed an aversive STM memory to 0.15 M sucrose. After 40 h of starvation, the mutants developed an appetitive STM. Numbers below box plots indicate pairs of reciprocally trained independent groups of male flies. The letter “a” marks a significant difference from random choice as determined by a one-sample sign test (*P* < 0.05). The Student’s t tests were used to determine differences between two groups. For differences between more than two groups one-way ANOVA with Tukey’s post hoc HSD test was used. (*P\** < 0.05; *P\*\** < 0.01; *P\*\*\** < 0.001).

To alter the internal energy status, flies were starved either for 16 h or for 40 h before the training (Figure 1). We used 3- to 5-day-old male flies to minimize differences in body weight and to control for differences in food preferences. We included *Tβh^nM18^* mutants lacking octopamine in the experiments, since they showed defects in energy metabolism and sucrose reward learning (Li et al., 2016; Schwaerzel et al., 2003). Flies that were starved for 16 h established a positive association between the odorant and the reward. They remembered this association for 2 min, but not for 6 h or 24 h. Consistent with previous results, learning and memory performance was significantly improved by prolonged starvation (Colomb et al., 2009). The memory was also remembered for longer, since it was still detectable 6 h after training. The *Tβh^nM18^* mutants starved for 16 h did not show 2 min memory but emerged memory 6 h later that lasted up to 24 h (Das et al., 2014; Schwaerzel et al., 2003). In contrast, *Tβh^nM18^* mutants remembered the rewarded odorant after only 2 min following starvation extended to 40 h. The memory was still detectable after 6 h and 24 h (Figure 1B). We repeated similar experiments using a lower concentration of 0.15 M sucrose as reinforcer (Figure 1C). In contrast to the experiments using 2 M sucrose as a reward, 16 h starved *Tβh^nM18^* mutants form a negative association with 0.15 M sucrose directly after the training, that turns into a positive association when the flies were starved longer 40 h. Again, prolonged starvation significantly increased learning and memory performance in controls and *Tβh^nM18^* mutants. Thus, the length of starvation and the reinforcer strength influence appetitive memory strength.

Next, we investigated how starvation affects the stability of memory. Protein synthesis-dependent long-term memory is still labile immediately after training and can be blocked by 4°C cold-shock anesthesia immediately after training. After consolidation, long-term memory is cold shock resistant and insensitive to a cold shock 1 h before the test (Krashes and Waddell, 2008). To investigate how hunger influences the stability of food-related memories, we administered cold shock anesthesia immediately after the training and 1 h before the test and analyzed the effect on memory formation in differently starved control and *Tβh^nM18^* flies (Figure 2A). We reduced the time for the memory test after training from 6 h to 3 h to be able to detect memory in 16 h starved flies. The memory was completely abolished by a cold shock directly after training or shortly before the test. Prolonging the starvation period resulted in memory that was sensitive to cold-shock directly after training but not shortly before the test. In contrast, 16 h starved *Tβh^nM18^* mutants developed memory that was sensitive to cold shock directly after training and cold shock insensitive shortly before the test. The results together with the observed memory 24 h later of *Tβh^nM18^* after the same training (Figure 1B) support that *Tβh^nM18^* mutant develop long-term memory. Longer periods of starvation in *Tβh^nM18^* mutants resulted in anesthesia-resistant long-term memory.

**Figure 2.**
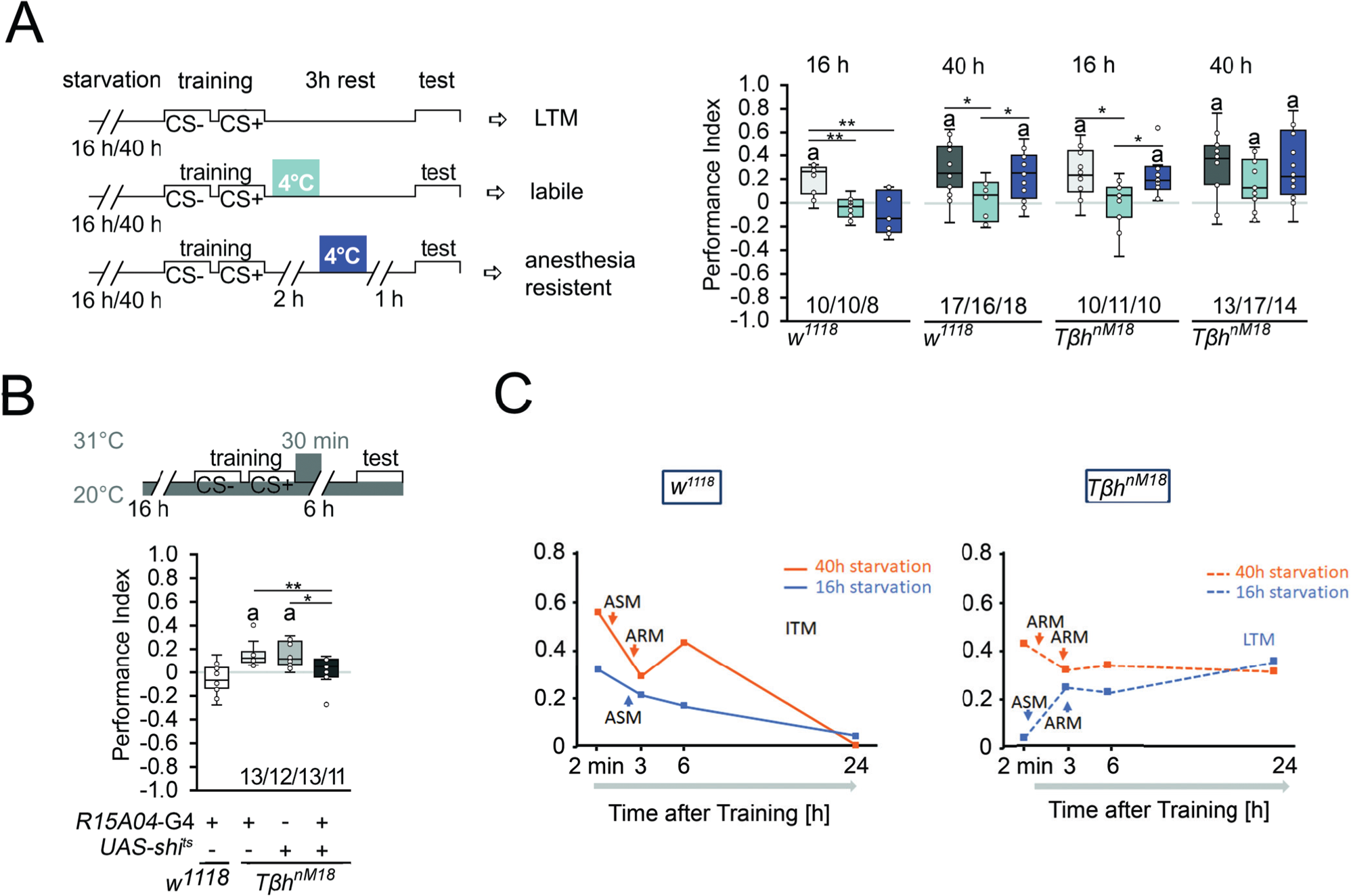
Starvation influences the type of memory. **A**, Appetitive STM training with 2 M sucrose and cold shock directly or 2 h after training. Mildly starved control flies exhibit an appetitive memory sensitive to cold shock 3 h after training. Severely starved control flies develop a memory that is initially sensitive to cold shock, but becomes insensitive after 2 h. This phenotype is shared with mildly starved *Tβh^nM18^* mutants. Prolonged starvation in the mutants shifts memory to cold shock insensitive memory. **B**, To block neuronal activity and the formation of LTM, a 30-min heat shift was applied immediately after training to flies expressing a temperature-sensitive *shibire* transgene under the control of the R1504-Gal4 driver. The block results in *Tβh^nM18^* mutants losing LTM. Numbers below box plots indicate pairs of reciprocally trained independent groups of male flies. The letter “a” marks a significant difference from random choice as determined by a one-sample sign test (*P* < 0.05). One-way ANOVA with Tukey’s post hoc HSD test or for data in **A**, and Kruskal-Wallis followed by post-hoc Dunn’s test and Bonferroni correction was used to determine differences in **B**. (*P\** < 0.05; *P\*\** < 0.01). **C**, Model summarizing memory performance of control and mutant flies, that were either starved for 16 h (blue line) or 40 h (orange line). The dots present the average of the data presented in Figure 1 and Figure 2. Anesthesia sensitive memory (ASM), anesthesia resistant memory (ARM), intermediate memory (ITM) and long-term memory (LTM).

To further confirm that the observed memory in 16 h starved *Tβh^nM18^* is indeed long-term memory, we wanted selectively inhibit long-term memory in *Tβh^nM18^* mutants and analyze their food-related memory (Figure 2B). We analyzed memory performance 6 h after training, as control flies show no memory at this time point (Figure 1B, Figure 2B). The R15A04-Gal4 driver targets dopaminergic neurons of the PAM cluster, which are specifically required for appetitive long-term memory but not short-term memory (Yamagata et al., 2015). Blocking the function of these dopaminergic neurons immediately after training using a temperature-sensitive *shibire* transgene (UAS-shi^ts^) and a 30-min heat pulse of 31°C resulted in loss of memory in *Tβh^nM18^* mutants (Figure 2B). Control flies carrying one copy of the Gal4 transgene showed no memory. Without heat shock, the *Tβh^nM18^* mutants carrying the UAS-shi^ts^, the R15A04-Gal4 or both transgenes developed memory (Figure S1). Thus, although they do not show memory immediately after training, *Tβh^nM18^* mutants that were briefly starved develop long-term memory.

These results, together with the course of memory loss, show that lightly hungry flies form a memory that lasts up to 3 h and is sensitive to anesthesia (Figure 2D). Since no memory can be observed 24 h later, this indicates that it is an anesthesia-sensitive intermediate memory (ITM). Prolonged starvation results in anesthesia-resistant ITM (Figure 2D). In contrast, mildly starved *Tβh^nM18^* mutants form a long-term memory that is sensitive to anesthesia-sensitive and detectable 24 h after the training. Prolonged starvation in *Tβh^nM18^* mutants leads to anesthesia-resistant long-term memory (Figure 2D).

Thus, the longer the starvation lasts, the more stable the memory becomes. Depending on the duration of starvation after the same training phase, the animals first form an STM memory, then anesthesia-sensitive or insensitive intermediate memory. Mildly starved *Tβh^nM18^* mutants do not form a short-term memory, but a long-term memory and after prolonged starvation, short-term and anesthesia resistant memory.

### Octopamine is a negative regulator of memory

Since the *Tβh^nM18^* mutants lack the neurotransmitter octopamine and form LTM, we next investigated whether octopamine normally suppresses LTM. To do so, we first blocked the function of octopamine receptors in controls immediately after the training by feeding the octopamine antagonist epinastin for 1 h and analyzed memory 5 h later (Figure 3A). If the octopamine receptor function is required for long-term memory, prolonged memory should also occur in control flies that normally show no memory after 16 hours of starvation. This was the case. Consistent with the idea that octopamine is a negative regulator of long-term memory, feeding octopamine to *Tβh^nM18^* mutants immediately after training blocked long-term memory (Figure 3A). To analyze whether octopamine is also able to block STM, we fed octopamine prior to training to control flies (Figure 3B). A short pulse of octopamine before the training inhibits STM.

**Figure 3.**
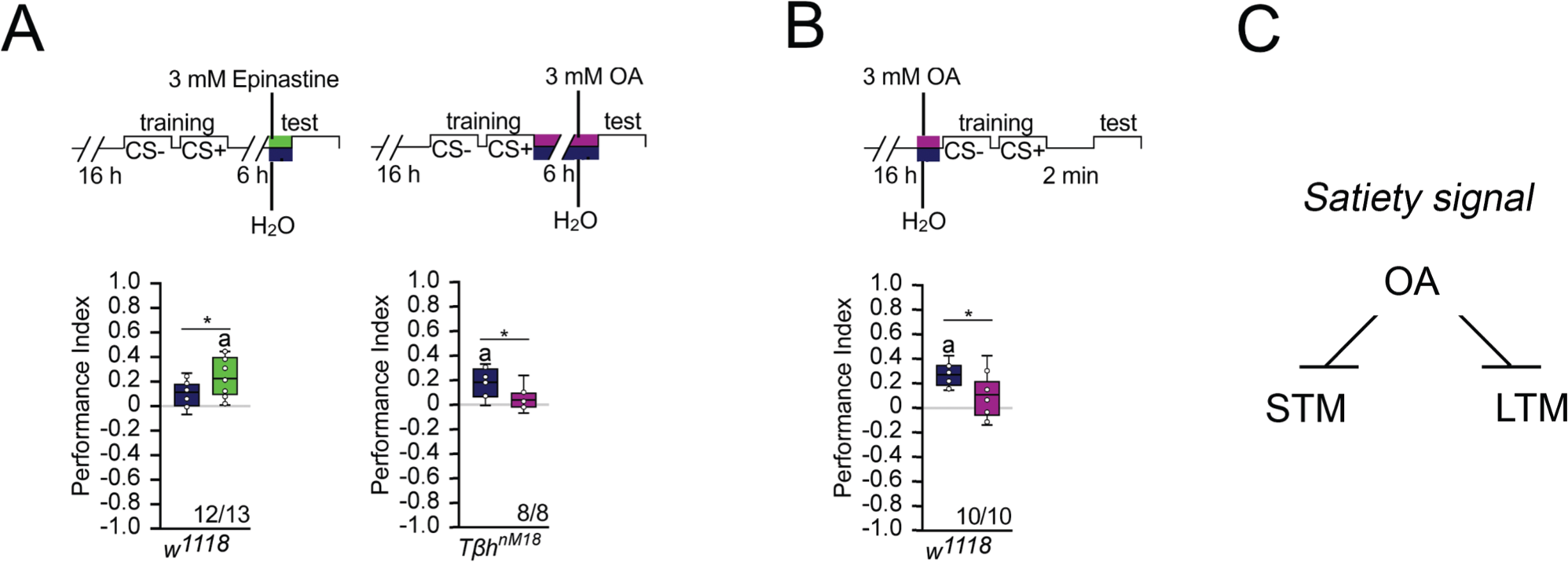
Octopamine suppresses memory. **A**, Feeding 3 mM of the octopamine receptor antagonist epinastine for 1 h after training resulted in memory 6 h later in *w^1118^* flies. Feeding 3 mM octopamine for 6 h after training suppresses LTM in *Tβh^nM18^*. **B**, A 3 mM octopamine feeding pulse 30 min before training inhibits STM in *Tβh^nM18^* mutants. Controls were water-fed. **A, B,** Flies were starved for 16 h and 2 M sucrose was used as reinforcer. The letter “a” marks a significant difference from random choice as determined by a one-sample sign test (*P* < 0.05). Student’s t tests were used to determine differences between two groups. (*P\** < 0.05). Numbers below box plots indicate pairs of reciprocally trained independent groups of male flies. **C**, Model for memory suppression.

We were wondering whether octopamine could reduce the appetite in starved animals and thereby reducing the value of the food reward. To investigate this, we fed starved flies with a short octopamine pulse and assayed their sucrose intake after 3 h (Figure S2). Flies fed with octopamine showed a significant reduction of sucrose intake, but not flies fed with a similar amount of tyramine or a 10-fold increase in tyramine concentration. Therefore, it is possible that increased octopamine content influences how food is evaluated. This is consistent with a function of octopamine as a signal for food or high internal energy levels. Taken together, octopamine is a negative regulator of appetitive long-term memory and STM (Figure 3C).

### Starvation influences sucrose consumption preference

Starvation reduces the internal energy storage. The reduction could lead to a reassessment of the external food cues and increased food consumption to restore the energy supplies. First, we investigated how starvation reduces the glycogen storage in whole animals (Figure 4A). In controls and *Tβh^nM18^* mutants, starvation reduces the glycogen level. However, non-starved *Tβh^nM18^* mutant males had significantly higher glycogen levels at baseline. After 40 h of starvation, the glycogen levels of *Tβh^nM18^* mutants were still higher than those of controls that were starved for the amount of time. Adult *Tβh^nM18^* mutants have a reduced sucrose intake (Li et al., 2016; Scheiner et al., 2014). The assessment of external food sources might be reflected in the choice of food. To examine whether hunger influences the evaluation of an external food stimulus, we starved flies and determined the preference for consuming sucrose over protein-enriched food using the capillary feeder assay (Figure 4B). Non-starved flies reared on standard fly food preferred sucrose over protein-enriched food, but *Tβh^nM18^* mutants to a significantly lesser extent than controls.18 h of starvation resulted in a decreased sucrose preference in controls and *Tβh^nM18^* mutants, but to a significantly lower level in *Tβh^nM18^* mutants. Extending the starvation period to 40 h resulted in a similar sucrose consumption in control and mutant flies.

**Figure 4.**
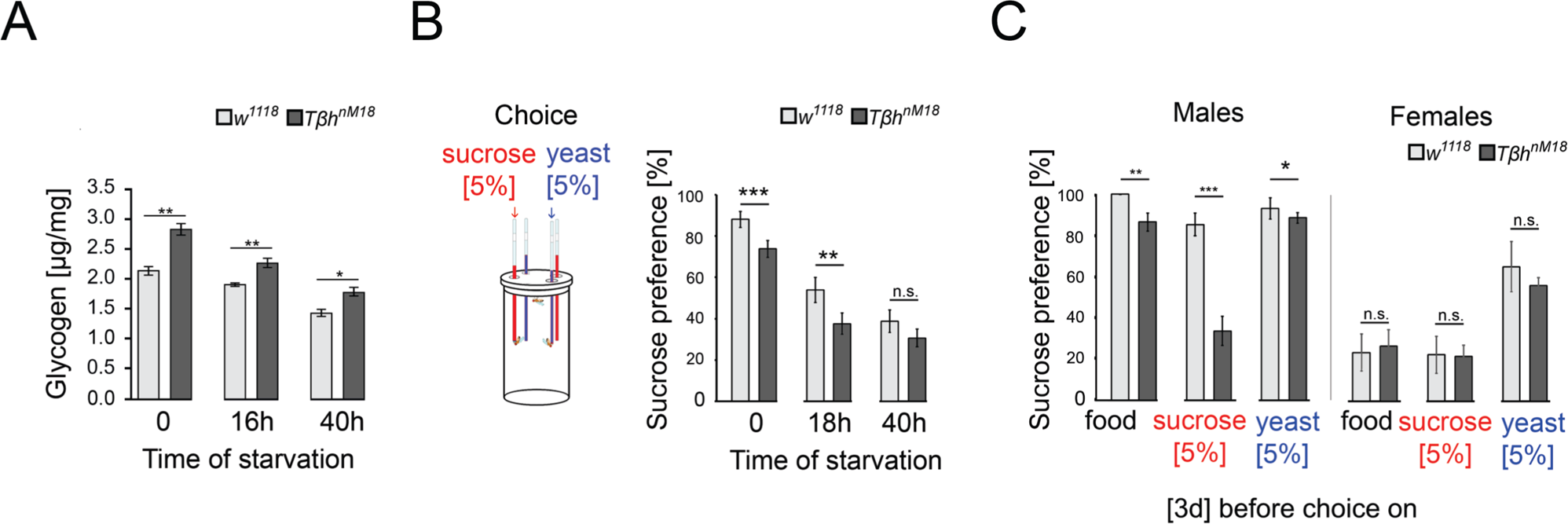
Elevated glycogen levels correlate with reduced sucrose preference. **A**, Analysis of whole-body glycogen levels in *w^1118^* and *Tβh^nm18^* flies. In *Tβh^nM18^* flies, glycogen content is significantly higher than in *w^1118^* flies under similar starvation conditions. N = 3 groups of 5 male flies. **B**, Flies were starved 18 h or 40 h before food intake was measured for 24 h. Flies chose between 5% sucrose and 5% yeast. The preference was determined. All flies showed a significant preference for sucrose consumption. Starvation reduced the preference. *Tβh^nm18^* showed a significantly reduced preference for sucrose in comparison to control flies, but not after 40 h starvation. N = 20 - 26 groups of eight flies. **C**, Feeding male flies for 3 days on standard fly food (food), 5% sucrose or 5% yeast resulted in control flies preferring to consume sucrose. Male *Tβh^nm18^* mutants fed normal food, sucrose and yeast showed a significant reduction in sucrose preference. Female flies of controls and mutants did not differ in their preferences. N = 14 - 28 groups of eight flies. To determine differences between two groups, the Mann-Whitney U test was used. *P\** < 0.05; *P\*\** < 0.01, *P\*\*\** < 0.001.

To further investigate whether *Tβh^nM18^* mutants integrate their internal glycogen storage into their feeding preference, we deprived male flies of sucrose- or protein-enriched food by feeding them 5% sucrose, 5% yeast or standard fly food for 3 days. The food preference was analyzed after deprivation (Figure 4C). As controls, we included mated females in our analysis as they have different nutritional requirements due to their mating status (Ribeiro and Dickson, 2010; Vargas et al., 2010). Male flies fed standard food for 3 days strongly preferred sucrose over protein-enriched food, with *Tβh^nM18^* mutants showing a significant reduction. Yeast-deprived and sucrose-deprived *Tβh^nM18^* mutant showed a significantly reduced preference for sucrose. In mated females, no differences were observed in food preference between controls and the *Tβh^nM18^* mutant. In conclusion, the reduced preference for sucrose consumption correlates in *Tβh^nM18^* with increased glycogen levels. Furthermore, *Tβh^nM18^* mutants can sense the reduction of specific internal energy stores and change their food preferences accordingly.

### Internal glycogen storage influences sucrose-related memories

Since memory performance increases when the internal energy supply is reduced by starvation, we wanted to investigate whether the internal energy supply influences memory performance. In *Drosophila*, glycogen is mainly found in the fat bodies - the major energy storage organ - and the muscles, a major site of energy expenditure (Wigglesworth, 1949). Glycogen synthase and glycogen phosphorylase control glycogen content in the body (Figure 5A). Knockdown of glycogen synthase using *GlyS^HMS01279^-RNAi* efficiently reduced glycogen levels in larvae, while knockdown of glycogen phosphorylase using *GlyP*^HMS00032^*-RNAi* efficiently increased glycogen levels (Yamada et al., 2018). We altered glycogen levels in the muscles using the *mef2*-Gal4 driver (Ranganayakulu et al., 1998) and/or the fat bodies using the FB-Gal4 driver (Gronke et al., 2003) and analyzed the effect of altered glycogen levels on short-term memory (Figure 5). We used PAS staining to confirm the down- or up-regulation of glycogen in larval muscles or fat bodies, respectively (Figure 5) (Yamada et al., 2018). In addition, we quantified the glycogen levels in the bodies of adult flies (Figure S3). As we could not always detect differences in glycogen content in whole flies, we also analyzed the glycogen level in the muscle-rich thorax or fat body-rich abdomen (Figure S3). The knockdown of glycogen phosphorylase increased glycogen levels and the knockdown of glycogen synthase reduced glycogen levels.

**Figure 5.**
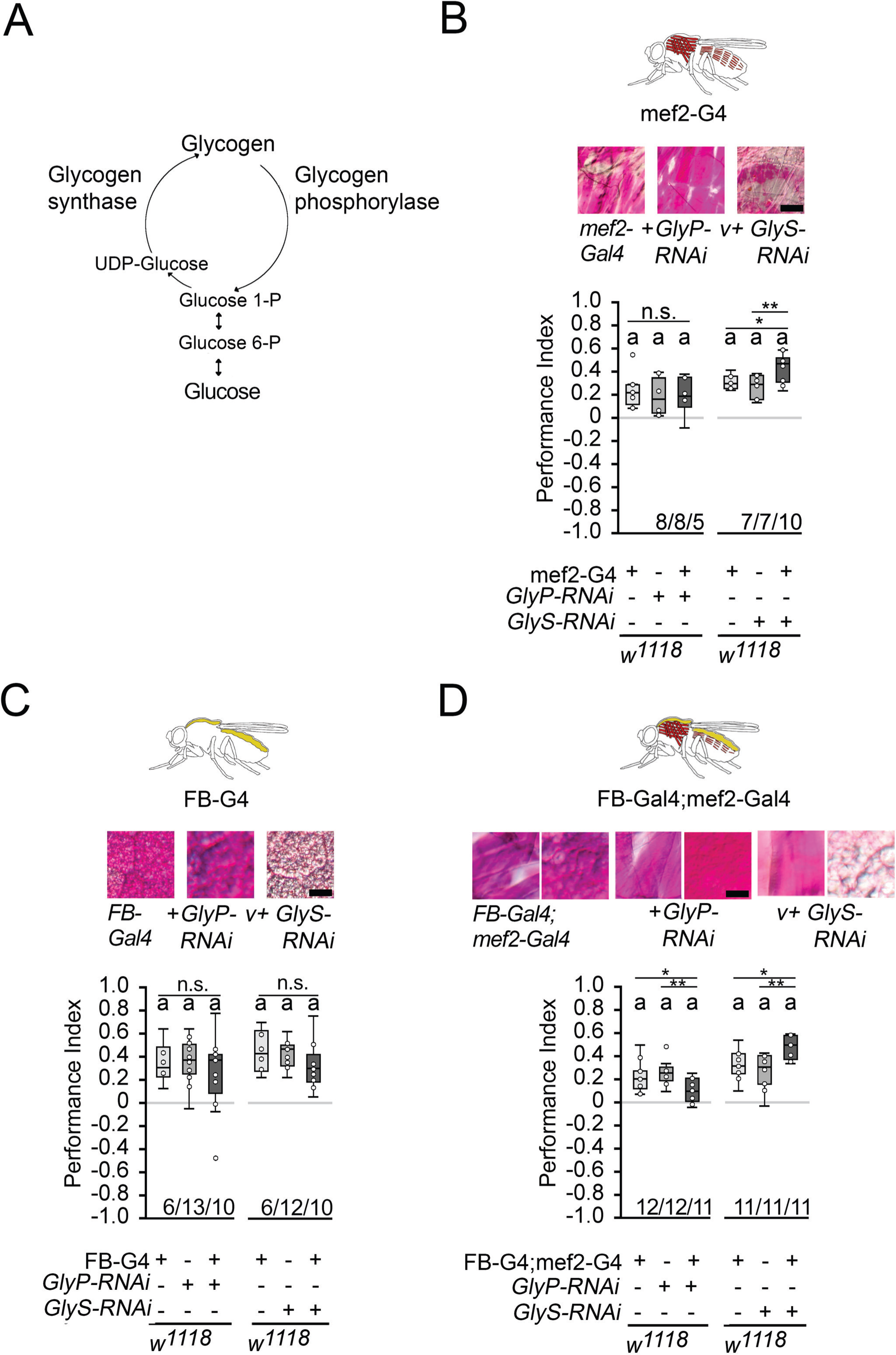
Carbohydrate storage influences appetitive STM. **A**, Schemata of glycogen synthesis. The expression of *GlyP-RNAi* reduced glycogen phosphorylase and increased glycogen levels, whereas *GlyS-RNAi* reduced glycogen synthase and decreased glycogen levels in target tissues (Figure S3). **B** - **D**, PAS was used to visualize glycogen levels in larval muscle or fat bodies. **B**, Increases in glycogen in the muscles have no effect on STM, whereas reduced muscle glycogen increases appetitive STM. **C**, Increased or decreased glycogen levels in the fat bodies did not interfere with STM. **D**, A combined increase in glycogen in muscles and fat bodies reduced STM, and a decrease in glycogen increased STM. Flies were starved for 16 h and 2 M sucrose was used as reinforcer. Numbers below box plots indicate one pair of reciprocally trained independent fly groups. The letter “a” marks a significant difference from random choice as determined by a one-sample sign test (*P* < 0.05). The one-way ANOVA with post hoc Tukey’s HSD was used to determine differences between three groups (*P\** < 0.05; *P\*\** < 0.01).

Increasing glycogen levels in the muscles had no effect on the short-term memory of the 16 h starved flies, but decreasing glycogen levels significantly improved memory performance (Figure 5B). Increasing or decreasing glycogen levels in the fat bodies had no effect on memory performance (Figure 5C). When the muscles and fat bodies glycogen levels were significantly increased, flies showed a reduced memory to odorants paired with sucrose. An increase in memory performance was observed when glycogen levels were significantly reduced in both tissues (Figure 5D). Recently, it has been shown that energy metabolism in mushroom bodies is important for the formation of LTM (Placais et al., 2017). To analyze the function of the mushroom bodies, the expression of *GlyS^HMS01279^-RNAi* under the control of the Mef2-Gal4 driver was repressed in the mushroom bodies using the mb247-Gal80 driver (Krashes et al., 2007). The memory was still increased (Figure S4). Conversely, the reduction of *GlyS^HMS01279^-RNAi* using the mb247-Gal4 driver targeting the mushroom bodies(Zars et al., 2000) did not alter short-term memory (Figure S4). Thus, low levels of glycogen in the muscles after starvation positively influence appetitive short-term memory, whereas high levels of glycogen in the muscles and fat body decrease reduce short-term memory.

### Internal glycogen levels reduce sucrose-related memories in Tβh mutants

The increased glycogen levels in *Tβh^nM18^* mutants could be responsible for the reduced STM. To determine whether the reduction in glycogen levels in the muscles or fat bodies restores STM, we expressed *GlyS^HMS01279^-RNAi* under the control of the *mef2*-Gal4 or *FB*-Gal4 driver in *Tβh^nM18^* mutants and analyzed STM (Figure 6A). Neither the reduction in the muscles nor the reduction in fat bodies of *Tβh^nM18^* mutants improved STM. Only when glycogen was reduced in both tissues did the *Tβh^nM18^* mutants show improved STM compared to controls. Thus, *Tβh^nM18^* mutant flies can form appetitive STM similar to controls when the energy storage is sufficiently reduced. Next, we analyzed whether male *Tβh^nM18^* mutants can form STM to other nutrients than carbohydrates by using a protein-enriched diet in the form of 5% yeast as a positive reinforcer (Figure 6B). To determine whether there is a difference in the evaluation of protein as a food source between male flies and *Tβh^nM18^* mutants, we measured yeast intake in non-starved and starved flies (Figure S5). Non-starved *Tβh^nM18^* mutant males have a significantly higher protein intake than controls. However, after 16 h of starvation, the protein intake was comparable to that of control flies. When 5% yeast was used as a food reward, male *Tβh^nM18^* mutants showed comparable STM levels controls (Figure 6B).

**Figure 6.**
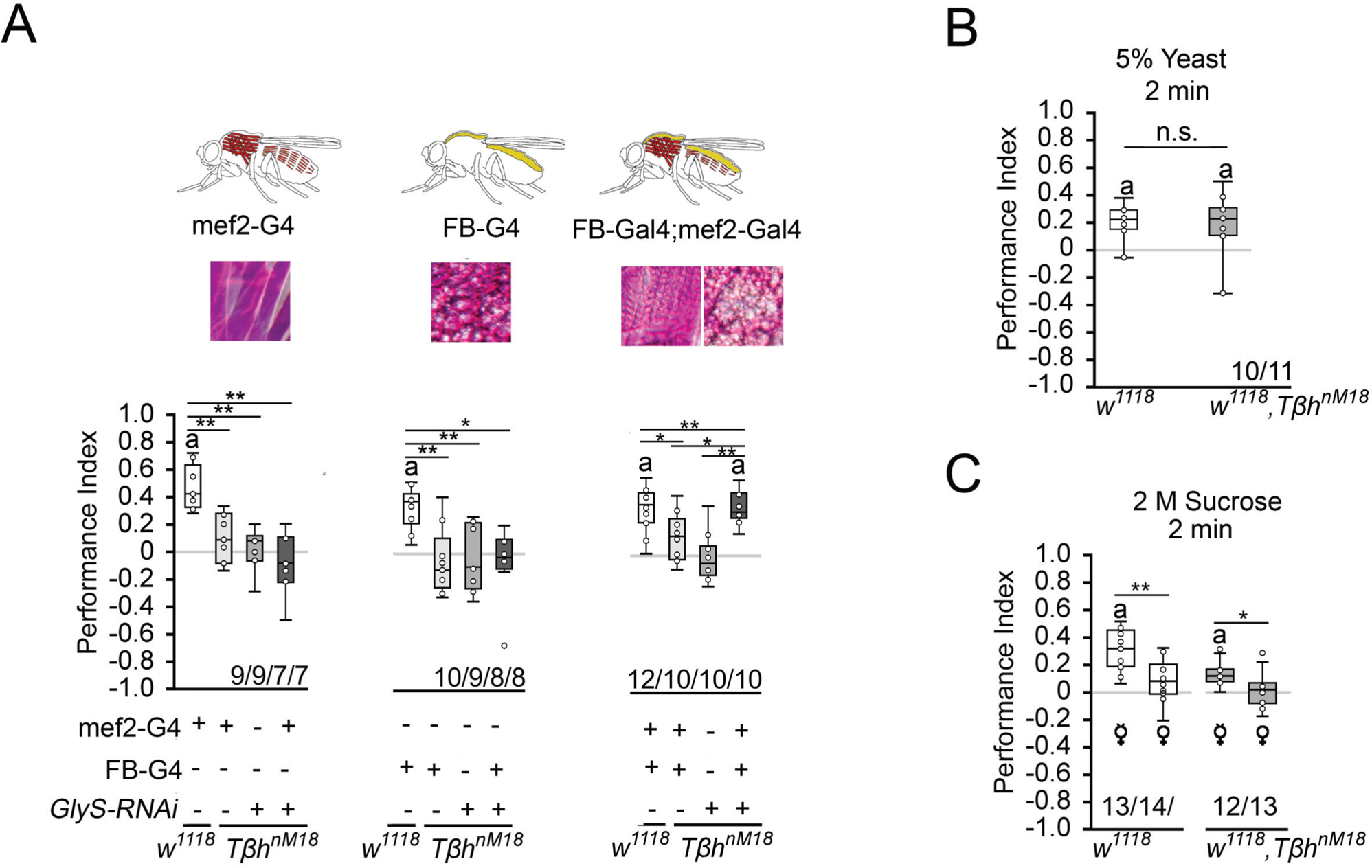
Reducing glycogen in *Tβh^nM18^* improves appetitive STM. **A**, Decreasing glycogen concentration using UAS-*GlyS^RNAi^* in the muscles or fat bodies in *Tβh^nM18^* mutants did not improve STM, but decreasing glycogen in both tissues improved STM to control levels. Flies were starved for 16 h and 2 M sucrose was used as reinforcer. **B**, 16 h starved *w^1118^* and *Tβh^nm18^* flies formed similar levels of appetitive STM when 5% yeast was used as a reinforcer. **C**, 16 h starved virgin females of w1118 and *Tβh^nm18^* displayed STM, whereas mated females of both genotypes did not. Differences from random choice were determined using a one-sample sign test and marked with the letter “a” (*P* < 0.05). Differences between two groups were determined using Student’s t tests, and differences among four groups were determined with one-way ANOVA with Tukey’s HSD post hoc test. *P\** < 0.05; *P\*\** < 0.01. Numbers below box plots indicate one pair of reciprocally trained independent fly groups.

To further analyze the influence of the internal energy status on memory performance, we took advantage of the observation that female flies have different nutritional requirements depending on their mating status (Ribeiro and Dickson, 2010; Vargas et al., 2010). Virgin females showed a higher consumption preference for sucrose than mated females when given the choice between yeast and sucrose (Figure 4C), demonstrating a difference in internal energy demands and suggesting a difference in valence for different diets. We tested virgin and mated female flies of controls and *Tβh^nM18^* mutants for STM with 2 M sucrose as reinforcer (Figure 6C). Virgin females remembered the odorant paired with sucrose significantly better than mated females. This was also true for *Tβh^nM18^* mutant females. Thus, the internal energy status influences how the reward is evaluated in learning and memory.

### Insulin-like signaling in octopaminergic neurons regulates STM

How is the internal energy status integrated into the reward system? Insulin-like signaling regulates glycogen levels in invertebrates and vertebrates (Semaniuk et al., 2021). Loss of the insulin receptor results in more circulating sugar but not increased glycogen levels (Shingleton et al., 2005). Thus, the insulin receptor might be a good candidate that links the internal energy level to reinforcing neurons. First, we analyzed whether the insulin receptor is expressed in octopaminergic reward neurons in the brain (Figure 7A). To detect the expression of an activated insulin receptor, we used an insulin antibody that recognizes the phosphorylated form of the insulin receptor (InR). This region is highly conserved between humans and flies. First, we tested whether the antibody indeed recognized the activated insulin receptor. Therefore, we overexpressed the activated insulin receptor using the UAS-InR.A1325D transgene under the control of the *dTdc2*-Gal4 driver (Figure S6). The InR.A1325D protein variant mimics the human V938D protein variant, which is constitutively active,(Longo et al., 1992) and the *dTdc2*-Gal4 driver targets octopaminergic reward neurons (Busch et al., 2009). The expression of activated InR resulted in increased immunoreactivity (Figure S6). The InR antibody therefore detects the activated InR. In the brain, activated InR is broadly expressed in a punctate manner and more specifically in the soma of *dTdc2*-Gal4-targeted neurons (Figure 7A). To uncouple energy sensing via the insulin receptor in reward neurons, we expressed UAS-InR.K1409A under the control of the *dTdc2*-Gal4 driver. The transgene encodes a dominant negative variant of the InR (InR^DN^) and disrupts with InR function (Wu et al., 2005). In animals starved for 16 h, the expression of InR^DN^ under the control of the *dTdc2*-Gal4 driver did not alter STM, but uncoupling of InR-dependent energy sensing in reward neurons in *Tβh^nM18^* restored STM to normal levels (Figure 7B and C).

**Figure 7.**
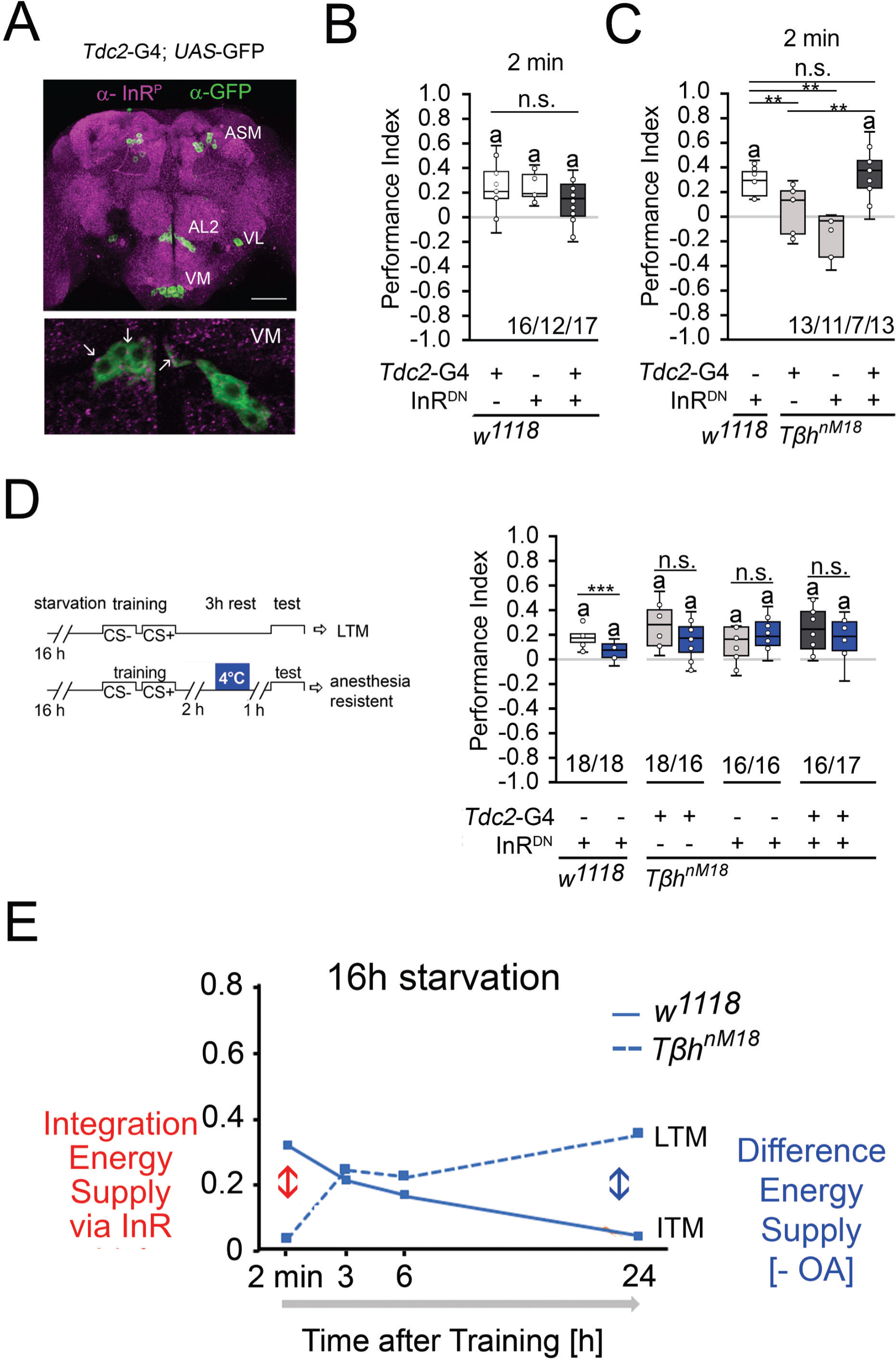
Insulin signaling in reward neurons regulates STM. **A**, The activated form of the InR is expressed in punctuate manner throughout the brain (in magenta) and is also detected in octopaminergic reward neurons visualized by using the UAS-mCD8::GFP transgene under the control of the *Tdc2*-Gal4 driver (in green). **B**, Blocking InR signaling in *Tdc2*-Gal4-targeted octopaminergic neurons does not change appetitive STM in 16 h starved flies using 2 M as reinforcer. **C**, Blocking InR signaling in *Tdc2*-Gal4-targeted octopaminergic neurons in *Tβh^nM18^* mutants restored STM to control levels. **D**, A cold shock did not disrupt emerging memory in *Tβh^nM18^* mutants with blocked InR under the control of the Tdc2-Gal4 driver. Student’s t test was used to determine differences between two groups, and one-way ANOVA with Tukey’s post hoc HSD test to determine differences between three or more groups. n.s. is not significant; *P\** < 0.05; *P\*\** < 0.01, *P\*\*\** < 0.01. The letter “a” marks a significant difference from random choice as determined by a one-sample sign test (*P* < 0.05). **E**, Model of memory regulation in *Tβh^nM18^* mutants. Long-term memory (LTM), intermediate-term memory (ITM).

To investigate whether the improved STM also affects the emerging LTM of the *Tβh^nM18^* mutants, we performed cold shock experiments in *Tβh^nM18^* mutants in which InR^DN^ was expressed in octopaminergic reward neurons (Figure 7D). The 3 h memory in controls is cold shock sensitive, but the memory in *Tβh^nM18^* mutants is cold shock insensitive, supporting the idea that the mutants formed anesthesia-resistant memory. Given that octopamine is a negative regulator of memory and that it is still absent in *Tβh^nM18^* mutants with blocked insulin signaling on octopaminergic reward neurons, it is not surprising that ARM is still observed in the mutants.

The results can be described with the following model (Figure 7E). Insulin receptor signaling on reward neurons integrated the internal energy status into memory formation. When energy levels are high enough due to increased glycogen levels or food intake, the insulin pathway suppresses the formation of STM. In return, the system releases octopamine to suppress the formation of food-related LTM. In mildly starved *Tβh^nM18^* mutants, the integration of the internal energy status into the reward neurons is still intact. In addition, the level of the internal energy supply combined with the loss of octopamine is still high enough to support the formation of protein biosynthesis-dependent LTM. In *Tβh^nM18^* mutants, two functionally different memory traces are formed after training, an insulin receptor-sensitive appetitive short-term memory and a long-term memory. In starved control animals, the glycogen levels are more severely depleted and octopamine is still present. This supports the formation of STM and an intermediate form of memory (ITM).

### Increased starvation results in overconsumption in Tβh^nM18^ mutants

To analyze whether the increased glycogen levels and the reduced sucrose reward of *Tβh^nM18^* mutants correlate with food consumption, the energy requirements in flies under different starvation conditions were analyzed by measuring food intake (Figure 8). Control flies starved for shorter or longer periods consumed similar amounts of 5% sucrose. In contrast, *Tβh^nM18^* mutants starved for 16 h consumed significantly less sucrose than controls. Longer-starved mutants consumed approximately 34% more sucrose (Figure 8B). After longer starvation, the glycogen levels in *Tβh^nM18^* mutants were still higher than those in controls (Figure 4A). Therefore, they overconsumed sucrose. The overconsumption was independent of the diet, as they showed similar overconsumption when fed with a solution containing 5% yeast and 5% sucrose (Figure 8B). To investigate whether the internal energy status is also integrated into the feeding behavior by the octopaminergic reward neurons, we blocked insulin signaling in reward neurons in 16 h-starved *Tβh^nM18^* mutants and analyzed sucrose consumption (Figure 8C). The reduced sucrose consumption of *Tβh^nM18^* mutants was significantly improved compared to controls. Thus, the octopaminergic reward system integrates via insulin receptor signaling the internal energy level into the regulation of food consumption. The mechanism is still intact in *Tβh^nM18^* mutants.

**Figure 8.**
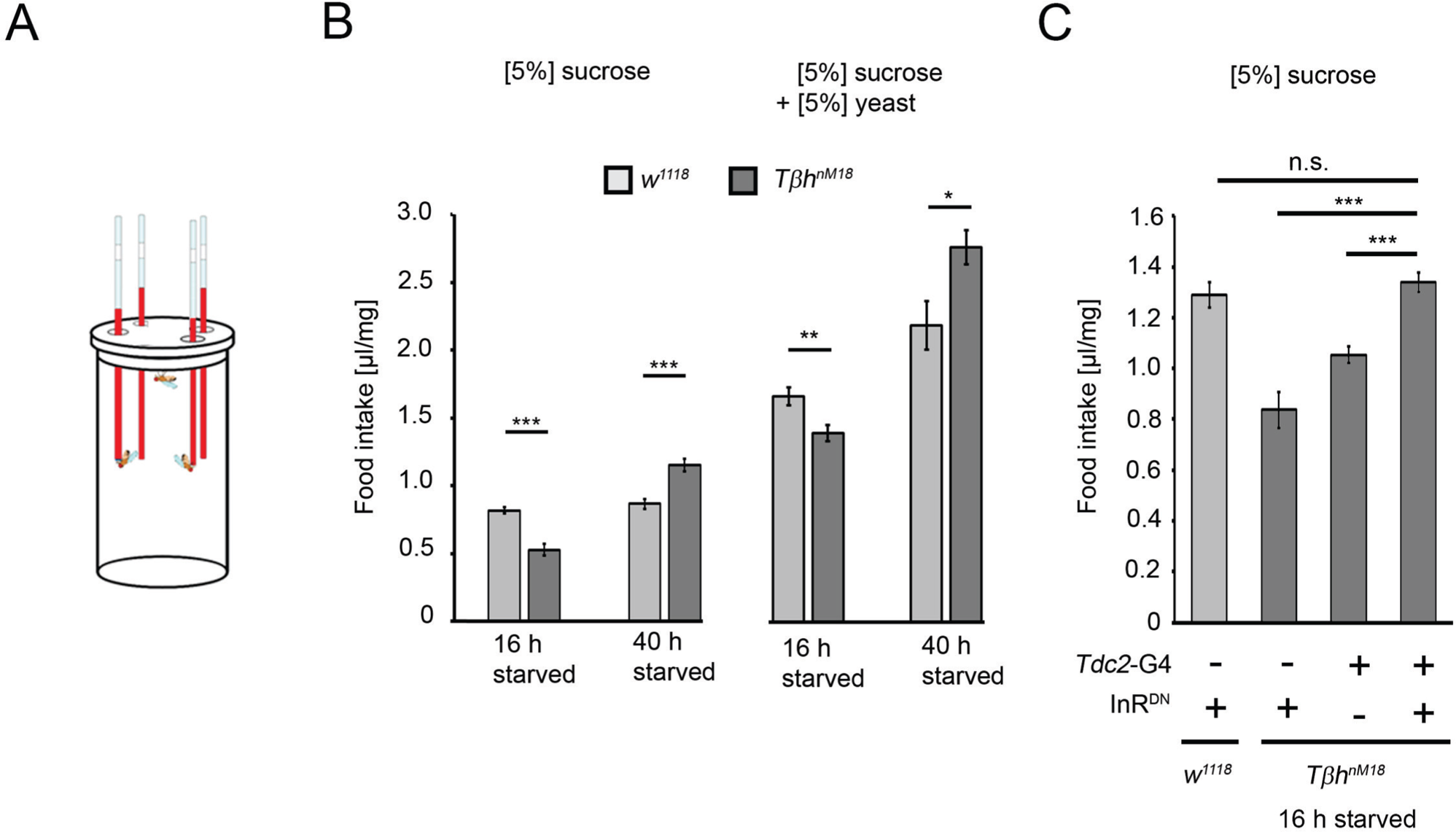
Prolonged starvation results in rebound sucrose intake in hyperglycemic *Tβh^nM18^* mutants. **A**, Capillary feeder assay used to determine food intake. **B**, Flies were starved 16 h or 40 h before 24 h food intake was measured. After 16 h of starvation, *Tβh^nM18^* mutants significantly consumed less 5% sucrose or 5% sucrose with 5% yeast. After 40 h of starvation, *Tβh^nM18^* mutants significantly consumed more sucrose and sucrose with yeast. N = 20 - 26 groups of eight flies. **C**, Blocking InR signaling in *Tdc2*-Gal4-targeted octopaminergic neurons in *Tβh^nM18^* mutants significantly increased 5% sucrose consumption. N = 20 - 28 groups of eight flies. To determine differences between two groups, Student’s t test was used, and to determine differences between three or more groups, one-way ANOVA with post hoc Tukey’s HSD was used. n.s. = non-significant; *P\** < 0.05; *P\*\** < 0.01, *P\*\*\** < 0.001.

## Discussion

Depending on the duration of starvation, flies develop STM and an anesthesia-sensitive intermediate type of a food-related memory. The extending the hunger period, memory formation shifts to an anesthesia-resistant intermediate memory. Presentation of the reward leads to the release of octopamine and this the suppression of LTM. The shifting of different memories phases in response to different energy supplies has been also described for memories that are formed of aversive non-food-related memories. Depending on the reduction of the internal energy supply, flies initially form protein-dependent aversive olfactory LTM and later ARM, a protein synthesis-independent form of memory (Wu et al., 2005). In times of energy shortage, the organism exchanges the costly protein biosynthesis-dependent LTM for the “less costly” protein biosynthesis-independent ARM (Mery and Kawecki, 2005). The effect of memory transition can also be observed at the level of appetitive LTM. In *Tβh^nM18^* mutants, mild starvation along with elevated glycogen levels leads to the formation of LTM and prolonged starvation leads to the formation of anesthesia-resistant LTM. Furthermore, *Tβh^nM18^* mutants form only anesthesia-sensitive aversive memory (Wu et al., 2013). In contrast, *Tβh^nM18^* mutants form cold shock-resistant LTM. Therefore, Tβh^nM18^ mutants are not defective in appetitive anesthesia-resistant types of memories rather the internal energy status defines how quickly anesthesia-resistant appetitive memories forms.

### Octopamine gates memory formation

What function does octopamine play in memory formation? Octopamine is a negative regulator of LTM, as feeding octopamine receptor antagonists to control flies results in memory, and emerging memory in *Tβh^nM18^* mutants is suppressed by octopamine after conditioning. Octopamine acts in the formation of LTM upstream of dopaminergic neurons, as blocking neurotransmitter release of dopaminergic LTM-mediating neurons immediately after conditioning blocks memory formation in *Tβh^nM18^* mutants. The results are consistent with the model describing that dopamine release is not required during the pairing of the conditioned stimulus and the reward rather than as gain control after the training (Adel and Griffith, 2021). The inhibitory function of octopamine controls the gain.

During the acquisition of appetitive STM, octopamine acts upstream of dopaminergic neurons M (Burke et al., 2012; Liu et al., 2012). However, at first glance, the function of octopamine appears to differ in STM and LTM. Loss of octopamine in *Tβh^nM18^* mutants results in loss of STM, indicating that octopamine is required as a positive regulator for appetitive STM (Schwaerzel et al., 2003). The loss of STM in *Tβh^nM18^* mutants has been attributed to the loss of labile STM which forms in response to the sweetness of sucrose, but not due to defects in stable LTM, which forms in response to the caloric value of sugar (Burke et al., 2012; Fujita and Tanimura, 2011). Sweetness is sensed by taste receptor that change their sensitivity during hunger (Inagaki et al., 2012; Marella et al., 2012) and thereby influencing the response to the external reinforcer. The responsiveness could be regulated by the octopaminergic system, as octopaminergic neurons regulate the sucrose sensitivity of gustatory neurons in *Drosophila* (Youn et al., 2018). However, the reduced sensitivity of gustatory receptors cannot explain why octopamine-deficient mutants do not form STM. The *Tβh^nM18^* mutants form aversive memory to lower concentrations of sucrose and the differences in sucrose preference of *Tβh^nM18^* do not translate to differences in food intake during training.

The internal energy status influences how the external source of sucrose is evaluated rather than octopamine being directly involved in the acquisition of appetitive STM. Consistently, *Tβh^nM18^* mutants can exhibit STM after prolonged periods of starvation. In mildly starved *Tβh^nM18^* flies that do not show STM, uncoupling of energy sensing in reward neurons by blocking insulin receptor function leads to STM. Furthermore, *Tβh^nM18^* mutant females with different energy requirements form appetitive STM, and a short pulse of octopamine before conditioning can even block STM. Thus, octopamine is not required in the acquisition of appetitive STM but rather suppresses LTM when enough energy is available. The function of octopamine as a suppressor of different memory types allows for selecting the memory type depending the internal and external information. This gating function in memory formation can also be observed in other behaviors. For example, octopamine regulates the decision to approach or turn away from a food source (Classen and Scholz, 2018). Taken together, the octopaminergic system integrates internal energy demands and the evaluation of external food supplies.

### The internal energy level of the animal influences memory formation

At the cellular level, insulin receptor signaling regulates energy metabolism (Chatterjee and Perrimon, 2021). The broad expression pattern of the insulin receptor in the *Drosophila* brain (Figure 7) indicates that every cell in the brain needs to regulate its own energy homeostasis. However, the fly’s entire energy resources and the evaluation of the external food supply must also be integrated. The reward system evaluates the external food supply. In mice, insulin receptor signaling in dopaminergic neurons mediates food reward (Konner et al., 2011). Consistently, insulin receptor function in octopaminergic reward neurons regulates the rewarding properties of sucrose in appetitive STM memory (Figure 6A) and during food intake (Figure 7C).

The systemic metabolic rate is regulated by fat bodies. For example, after food intake, the fat body secretes Unpaired 2 (Upd2), which in turn regulates the secretion of Drosophila insulin-like peptides via GABAergic neurons (Rajan and Perrimon, 2012). We show that the reduction of glycogen in the fat bodies is not sufficient to change appetitive STM, but reduction in the muscles or both tissues is. The results support a feedback mechanism between the assessment of glycogen levels in the muscles and the brain. In addition, the muscles and the fat bodies communicate about the energy content of both structures, and this information is signaled back to the brain. Such long-range signals from the muscles exist. Skeletal muscles are secretory organs (Pedersen and Febbraio, 2012). For example, the muscle-secreted Amyrel amylase reduces the age-related accumulation of polyubiquitinated proteins in the brain (Rai et al., 2021). Regardless of how the glycogen levels of both tissues are communicated to the brain, they influence how the reinforcer is evaluated. The feedback between the energy levels in both structures is still intact in octopamine-deficient mutants, as *Tβh^nM18^* mutants still form STM when the energy storage is sufficiently reduced.

### Are food-related memories and internal glycogen levels predictive of food intake?

The feeding behavior in hungry animals is regulated by various neural systems, including networks that receive and process sensory information and networks that assign reward properties to food (Berridge, 2009). Blocking insulin receptor function in dopaminergic neurons results in increased food intake and weight gain in mice (Konner et al., 2011). Similarly, blocking insulin receptor function in octopaminergic reward neurons in flies with increased glycogen levels and reduced food intake increases food intake. Thus, the octopaminergic system assigns reward properties to food. However, blocking insulin receptor signaling on reward neurons does not result in extensive overconsumption, supporting that the regulation of the food quantity is still intact and is not due to insulin receptor resistance in reward neurons. Increased levels of internal energy, e.g., glycogen, reduced the reward properties of food, resulting in a decreased positive association of food-related memories (Figure 5D; Figure 7A) and decreased food consumption (Figure 8). This is intended to prevent increased food intake when there is excess food. Insulin resistance could contribute to weight gain, because in addition to high internal levels of glycogen, the flies show normal food intake. However, the overconsumption only becomes visible after a prolonged period of hunger, which correlates with the emerging stable appetitive long-term memory. This suggests that the stability of appetitive memory could lead to the re-evaluation of food and could trigger the overconsumption of food.

In summary, the octopaminergic neurotransmitter system integrates several aspects that influence the regulation of food memories. Octopamine modulates the sensory perception of the conditioned stimulus and the sensory perception of the reward. Here, we show that the evaluation of food reward in the context of the energy storage is integrated by the octopaminergic system and influences the stability of food-related memories. The function of octopamine as a negative regulator of various forms of memories allows for selective regulation of food related behaviors such as food intake, and the loss of this regulation could also promote dysregulation of food intake. The close relationship between the octopaminergic neurotransmitter and the noradrenergic neurotransmitter system suggests that the integrator function may be conserved.

## Material and methods

### Drosophila melanogaster

Flies were raised on an ethanol-free standard cornmeal-molasses food at 25°C and 60% relative humidity on a 12 h/12 h day-night cycle. The following lines were used: *w^1118^*; *w^1118^,Tβh^nM18^*(Monastirioti et al., 1996); *UAS-GlyS^HMS01279^-RNAi* (BDSC #34930); *UAS-GlyP*^HMS00032^*-RNAi* (BDSC #33634)*; w^1118^*; *UAS-shi^ts^*(Kitamoto, 2001); w^1118^; P{UAS-InR. K1409A} (BDSC#8253); *w^1118^;FB-Gal4* (a generous gift from the Partridge Lab); *mef2-Gal4* (Ranganayakulu et al., 1998); R15A04-Gal4 (BDSC #48671); *Tdc2-Gal4* (Cole et al., 2005). All lines were backcrossed to *w^1118^* (Scholz Lab) for at least five generations to isogenize the genetic background. For behavioral experiments, three- to five-day-old male flies were used, if not otherwise indicated. Male flies of the F1 generation carrying one copy of the transgene were used as controls for behavioral experiments. Animal studies using *Drosophila melanogaster* were conducted in agreement with the regulations of the DFG and the Land North Rhine-Westphalia. Other ethics approval and informed consent statement are not applicable for research using *Drosophila*.

### Olfactory learning and memory

Associative olfactory learning and memory was trained and tested with a modified version of the Tully and Quinn olfactory conditioning apparatus (Schwaerzel et al., 2003; Tully and Quinn, 1985). Approximately 70 one- to two-day-old male flies were collected with CO_2_ anesthesia and were kept for 2 days at 25°C to recover from CO_2_ sedation. Briefly, three- to five-day-old male flies were starved for 16 h or 40 h in vials with water-soaked filter paper at the bottom. Flies were transferred to training tubes and exposed to the first odorant for 2 min, either 3-octanol (3-OCT diluted 1:80 in paraffin oil) or 4-methylcyclohexanol (MCH diluted 1:100 in paraffin oil). After that, they were transferred to a second tube and exposed to the second odorant in the presence of filter paper soaked with either 0.15 M sucrose, 2 M sucrose or 5% yeast. To analyze the abilities of the flies to learn and remember the odorant paired with the reward, 2 min, 3 h, 6 h or 24 h after training, flies were given the choice between odorant 1 (CS+) and odorant 2 (CS-) for 2 min. The performance index (PI) was calculated as PI = (# (CS+) + # (CS-) / (total # flies), where CS+ indicates the odorant associated with the appetitive reinforcer and CS-indicates the odorant that was not associated with the reinforcer. To exclude non-associative effects, each “n” consists of a reciprocally trained, independent group of naïve flies. For pharmacological experiments, flies were fed water, 3 mM OA or 2 M sucrose before training or with 3 mM OA or 3 mM epinastine between training and testing. The assay was performed at RT with 60% relative humidity. For cold shock experiments, flies in vials were placed in ice-cold water for 2 min.

Odorant acuity and odorant balance were tested with naïve flies that were not used later for learning and memory experiments. For testing, the flies were placed into the Tully Quinn setup. They chose between the odorant- or the solvent paraffin oil-containing sides or between both odorants for 2 min. The sucrose and yeast preference was determined by analyzing whether they preferred the side with a sucrose or yeast-soaked filter paper over a water-soaked filter paper. To determine sucrose intake, flies were fed with a food-safe colored sucrose solution and food intake was analyzed by the color in the abdomen (Table S1).

### Food intake

To analyze the consumption of nutrients in flies, the capillary feeder (CaFe) assay was performed (Diegelmann et al., 2017; (Ja et al., 2007)). Briefly, eight three- to five-day-old male or female flies that were either non-starved or starved for 18 h or 40 h had access to 4 capillaries filled with either 5% sucrose or 5% yeast or a mixture of both for 24 h at 25°C and 60% humidity. The solution was colored with red food dye. During starvation, flies were kept in vials with wet filter paper at 25°C and 60% humidity. The amount of consumed solution was determined using an electronic caliper. To account for evaporation, the average evaporation of three CAFE setups without flies was measured and used to normalize the average food intake per fly. To normalize food intake to body weight, at least 5 times 100 male flies per genotype and condition were weighed, and the average body weight of a single fly was determined. The total consumption per µg fly was calculated by dividing the total consumption per fly by the mean weight of a fly. N indicates the number of tested groups.

### Glycogen content

Whole body glycogen levels were determined with the Glucose (HK) Assay Kit (Sigma Aldrich, #GAHK20-1KT) according to the protocol of (Tennessen et al., 2014). A group of five male flies that were either sated or starved for 16 h or 40 h were homogenized in 100 µl ice-cold 1x PBS. To reduce enzymatic degradation, proteins and enzymes were heat inactivated at 70°C for 10 min. The supernatant was removed and diluted 1:3 with 1x PBS. Twenty microliters of the samples were either added to 20 µl of 1x PBS or PBS/Amyloglucosidase mix and incubated at 37°C for 60 min. Then, 100 µl of HK-reagent was added to the sample or glucose standard and incubated for 15 min at RT. Absorbance was measured at 340 nm. The glycogen content was calculated by subtracting the total glucose concentration from 1x PBS-treated samples from the total glucose of amyloglucosidase-treated samples.

### Periodic acid staining

To visualize glycogen levels in the fat bodies and muscle tissue of larvae and adult *Drosophila*, PAS staining was performed after Yamada et al., 2018 (Yamada et al., 2018) with a slight modification. The samples were fixed in 3.5% formaldehyde for 20 min and washed two times for 5 min with 1% BSA/PBS. Periodic acid solution was added for 5 min, followed by two washes of 5 min with 1% BSA/PBS. Schiff’s Reagent was added for 5 min, followed by two washes of 5 min with 1% BSA/PBS. Tissue was stored in 50% glycerol.

### Immunohistochemistry

For immunohistochemistry, antibodies raised in rabbits against the activated insulin-like receptor (cell signaling technology #3021) were used at a dilution of 1:50 in 5% normal goat serum in PBS with 0.1% Triton and incubated for two days. The brains were washed with PBS with 0.3% Triton.

## Quantification and statistical analysis

Food intake was displayed as the mean ± s.e.m. For learning and memory experiments, the data were displayed as boxplot ± minimum (Q1–1.5*IQR) and maximum (Q1 + 1.5*IQR). We used the Shapiro-Wilk test (significance level *P* < 0.05) followed by a QQ-Plot chart analysis to determine whether the data were normal distributed. For normal distributed data, we used the Student’s t test to compare differences between two groups and the one-way ANOVA with Tukey’s post hoc HSD test for differences between more than two groups. For nonparametric data, we used the Mann-Whitney U test for differences between two groups and for more than two groups the Kruskal-Wallis test with post hoc Duenn analysis and Bonferroni correction. The nonparametric one-sample sign test was used to analyze whether behavior was not based on random choice and differed from zero (*P* < 0.05). The statistical data analysis was performed using statskingdom (https://www.statskingdom.com).Boxplots were generated with Microsoft Excel 2016 and GIMP 2.10.12.

## Data availability

All data related to figures are included in the supplement Table S2.

## Acknowledgments

We thank the Linda Partridge lab and the Bloomington *Drosophila* Stock Center for providing *Drosophila* fly lines. H.S. was supported by SFB1340, DFG-Scho656-10-1 and the Hetzler award.

## Author contributions

H.S. initiated the project. M.B., H.S. and K.D. designed the experiments. M.B., K.A., K. D., M.F. and T.E.K. performed the experiments and, together with H.S., analyzed the data. M.B, M.F. and K.A. performed learning and memory tests. K.D. characterized insulin receptor expression. M.B. and T.E.K. performed food consumption assays. M.B. wrote a draft of the manuscript, and H.S. wrote the manuscript.

## Supplemental information titles and legends

Supplemental information includes 6 figures and two tables and can be found with this article online.

**Figure S1 Controls for heat shift experiment in Figure 2B**

To control for memory performance without a heat-shift, flies expressing a temperature-sensitive *shibire* transgene under the control of the R1504-Gal4 driver were trained and tested at the permissive temperature and compared to the respective controls. The tested groups showed memory 6 h after the training and did not differ from each other. Flies were starved for 16 h and 2 M sucrose was used as reinforcer. The behavior differed from random choice as determined using a one-sample sign test and marked with the letter “a” (*P* < 0.05). The one-way ANOVA with post hoc Tukey’s HSD was used to analyze difference between groups. n.s = no significant differences.

**Figure S2. Octopamine but not tyramine suppresses food intake (related to Figure 3)**

The *w^1118^* flies were starved for 17 h followed by feeding 1 h octopamine [10 mg/ml], tyramine [10 or 100 mg/ml] or water. The intake of 5% sucrose after 3 h was determined using the CAFÉ assay. Student’s t tests were used to determine differences between two groups and one-way ANOVA with Tukey’s post hoc HSD test between three groups. (*P\*\*\** < 0.001; n.s. = non-significant). N = 13 – 20 groups of 20 male flies.

**Figure S3. Glycogen level in adult flies with reduced GlyP and GlyS (related to Figure 5)**

The relative glycogen content in flies with altered GlyP and GlyS in muscles, fat bodies or both. The expression of *GlyP-RNAi* and *GlyS-RNAi* under the control of *mef2*-Gal4 did not significantly change the relative glycogen level in whole animals, but in the thorax. The expression of *GlyP-RNAi* under the control of *FB*-Gal4 increased significantly glycogen levels in whole animals, and the expression of *GlyS-RNAi* decreased glycogen in the abdomen. When both drivers are combined, the expression of *GlyP-RNAi* increases glycogen in whole animals and the expression of *GlyS-RNAi* decreases the glycogen levels. Glycogen was measured in 3 groups of 5 male flies and normalized to the protein levels and to the glycogen levels of flies of the Gal4 driver. ANOVA with post hoc Tukey’s HSD was used to determine differences between three groups. (*P\** < 0.05; *P\*\**< 0.01, *P\*\*\**< 0.001).

**Figure S4. The glycogen level in the mushroom bodies does not influence appetitive STM (related to Figure 5)**

**A**, The expression of *GlyS-RNAi* under control of the mef2-Gal4; mb247-Gal4 drivers increased STM. **B**, The expression of *GlyS-RNAi* under control of the mb247-Gal4 driver did not change STM. Flies were starved for 16 h and 2 M sucrose was used as reinforcer. One-way ANOVA with Tukey’s HSD post hoc test was used to determine differences between the groups. (*P\**< 0.05; *P\*\** < 0.01; n.s. = non-significant). The letter “a” marks a significant difference from random choice as determined by a one-sample sign test (*P* < 0.05). Numbers below box plots indicate one pair of reciprocally trained independent fly groups. **C**, Sensory acuity tests for the genotypes used.

**Figure S5. Starvation influences yeast consumption of *Tβh^nm18^* flies (related to Figure 6)**

Non-starved *Tβh^nm18^* male flies consumed significantly more yeast within 24 h. Starved *Tβh^nm18^* mutants consumed similar amounts of yeast. N = 26 groups of eight flies. To determine differences between both groups, Student’s t test was used. \*\*\**P* < 0.001; n.s. = non-significant difference.

**Figure S6. The antibody against phosphorylated InR recognizes activated InR (related to Figure 7)**

**A**, **B**, In the antennal lobes (AL) of the male adult brain, the immunoreactivity recognized by the anti-InR^P^ antibody is shown in magenta, and GFP expression is shown in green. The Gal4 expression of the *Tdc2*-Gal4 driver targeting octopaminergic neurons is visualized using the UAS-mCD8::GFP transgene. **B**, The expression of the constitutively active insulin receptor results in increased immunoreactivity detected by the anti-InR^P^ antibody.

**Table S1. Sensory acuity**

Odorant acuity and odorant balance were tested in independent fly groups of each genotype used in the behavioral experiments. For testing, flies were placed in the Tully Quinn setup. To analyze odorant preference and balance, they were allowed to choose between the odorant- or paraffin oil-containing sides or between both odorants for 2 min. Preference for sucrose and yeast was determined by analyzing whether flies preferred the side with a sucrose- or yeast-soaked filter paper over a water-soaked filter paper. Sucrose intake was analyzed by examining whether flies absorbed colored sucrose solution into the abdomen.

**Table S2. Data related to Figures**

