## Supplemental Figures S1-S6 for "Octopamine integrates the status of internal energy supply into the formation of food-related memories"

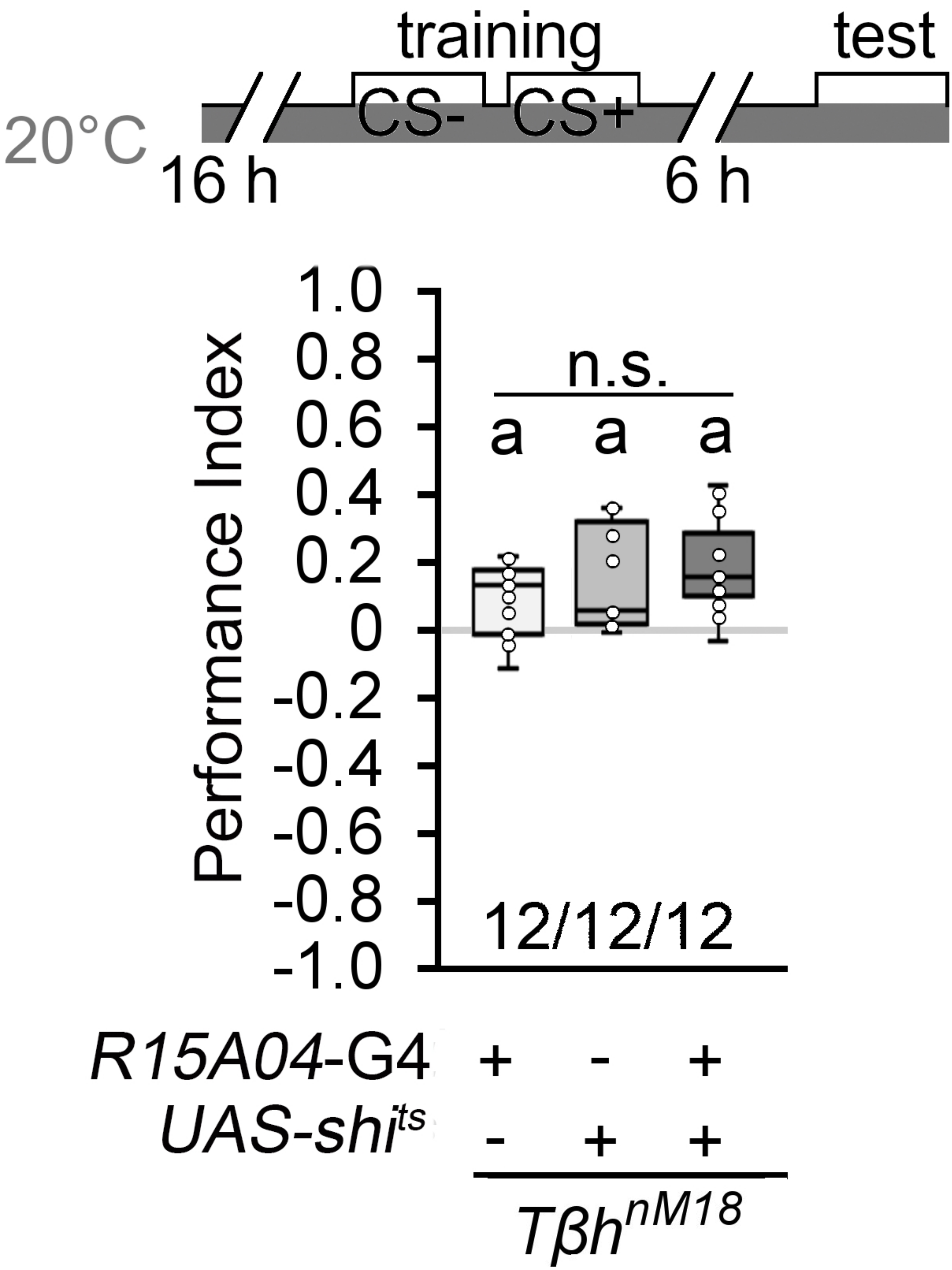
training
test
CS-
CS+
20°C
16 h
6 h
Performance Index
1.0
0.8
0.6
0.4
0.2
0
-0.2
-0.4
-0.6
-0.8
-1.0
n.s.
a a a
12/12/12
R15A04-G4 + - +
UAS-shits - + +
Tβh nM18

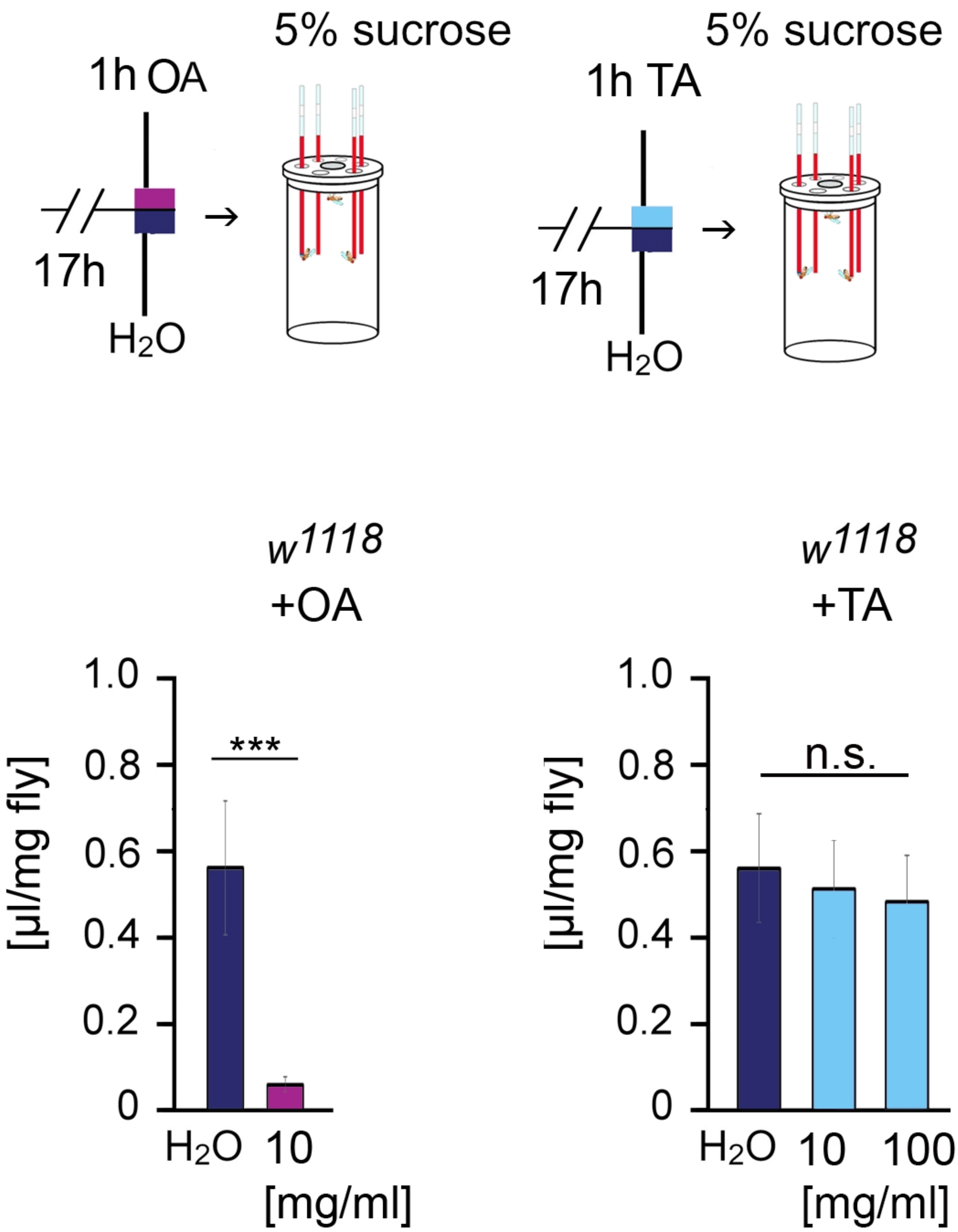

5% sucrose
1h OA
17h
H2O
5% sucrose
1h TA
17h
H2O
w1118
+OA
w1118
+TA
[µl/mg fly]
1.0
0.8
0.6
0.4
0.2
0
***
H2O
10
[mg/ml]
n.s.
H2O
10
100
[mg/ml]

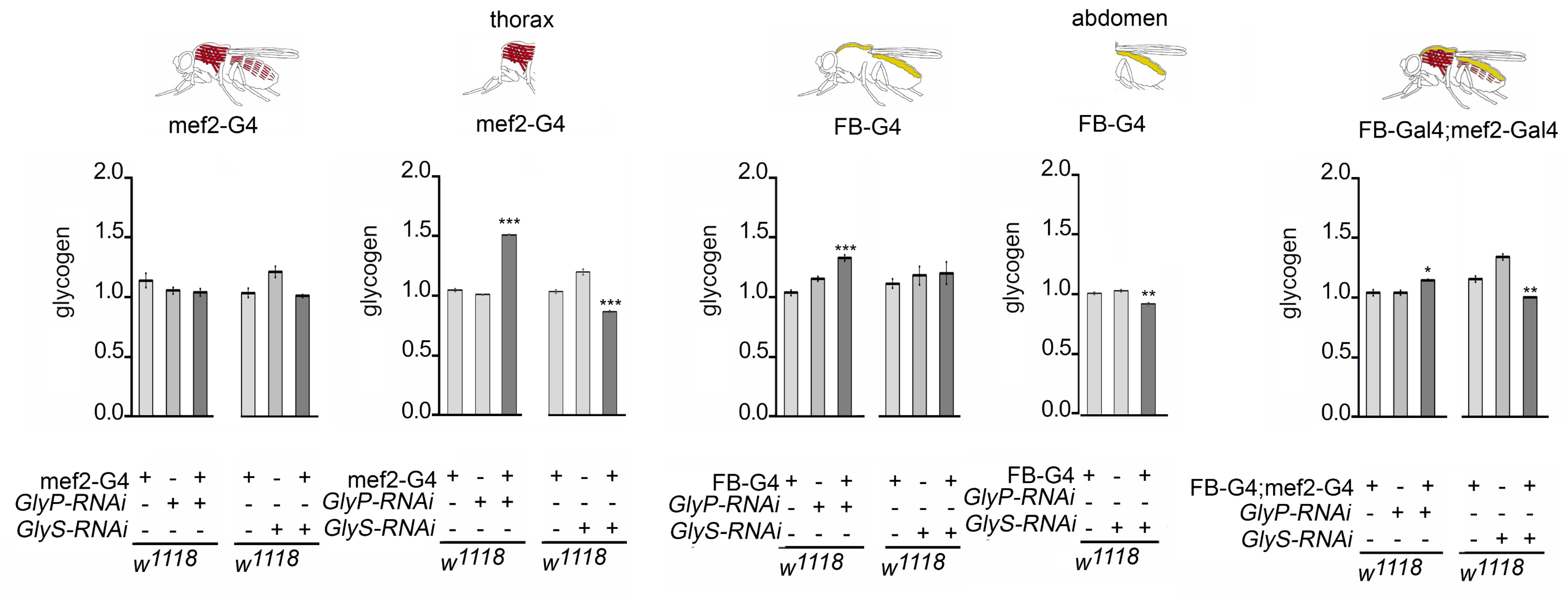
thorax
abdomen
mef2-G4
mef2-G4
FB-G4
FB-G4
FB-Gal4;mef2-Gal4
glycogen
2.0
1.5
1.0
0.5
0.0
***
*
**
mef2-G4
GlyP-RNAi
GlyS-RNAi
FB-G4
FB-G4;mef2-G4
w1118

A B

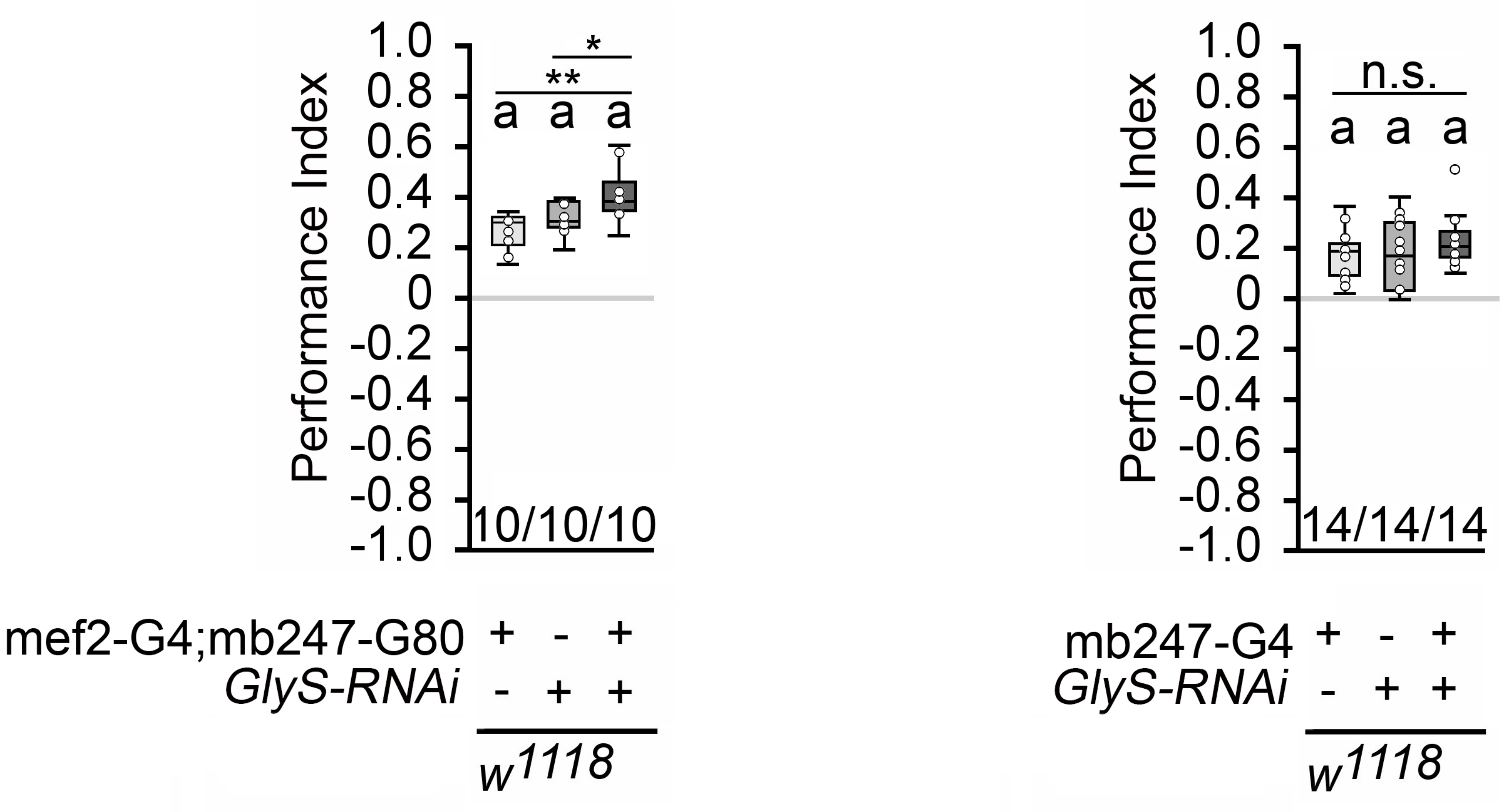


C

Sensory acuity

| Genotype | 3-Oct | MCH | Balance | Sucrose preference | N |
|---|---|---|---|---|---|
| mef2-G4; mb247-G80 | - 0.61 ± 0.07* | - 0.61*± 0.08 | 0.1 ± 0.06 | 0.3± 0.1* | 8 - 11 |
| UAS-GlyS RNAi/+ | - 0.53 ± 0.07* | - 0.52* ± 0.07 | 0.05 ± 0.07 | 0.22 ± 0.05* | 8 - 11 |
| mef2-G4; mb247-G80; UAS-GlyS RNAi | - 0.48 ± 0.07* | - 0.45* ± 0.07 | 0.1 ± 0.05 | 0.32 ± 0.08* | 8 - 11 |

| | 3-Oct | MCH | Balance | Sucrose preference | N |
|---|---|---|---|---|---|
| mb247-G4/+ | - 0.31 ± 0.08* | - 0.18* ± 0.03 | - 0.13 ± 0.4 | 0.15± 0.03* | 8 - 10 |
| UAS-GlyS RNAi/+ | - 0.37 ± 0.08* | - 0.29* ± 0.05 | 0.11 ± 0.16 | 0.29 ± 0.05* | 8 - 10 |
| mb247;UAS-GlyS RNAi | - 0.48 ± 0.1* | - 0.57* ± 0.0 | - 0.09 ± 0.24 | 0.28 ± 0.07* | 8 - 10 |

*P** < 0.05 significant difference from random choice

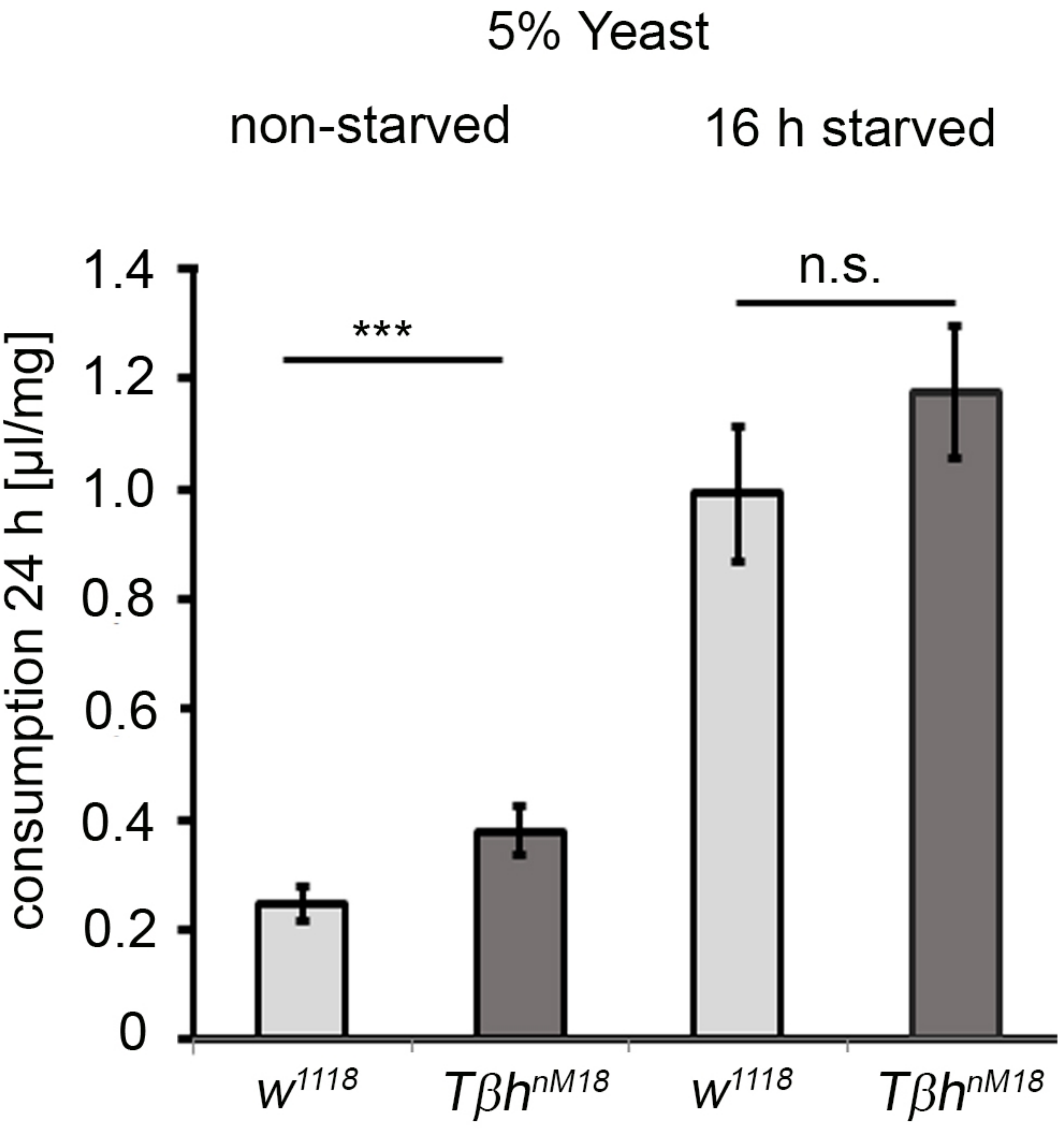

5% Yeast
non-starved
16 h starved
consumption 24 h [µl/mg]
1.4
1.2
1.0
0.8
0.6
0.4
0.2
0
***
n.s.
w1118
Tβh nM18
w1118
Tβh nM18

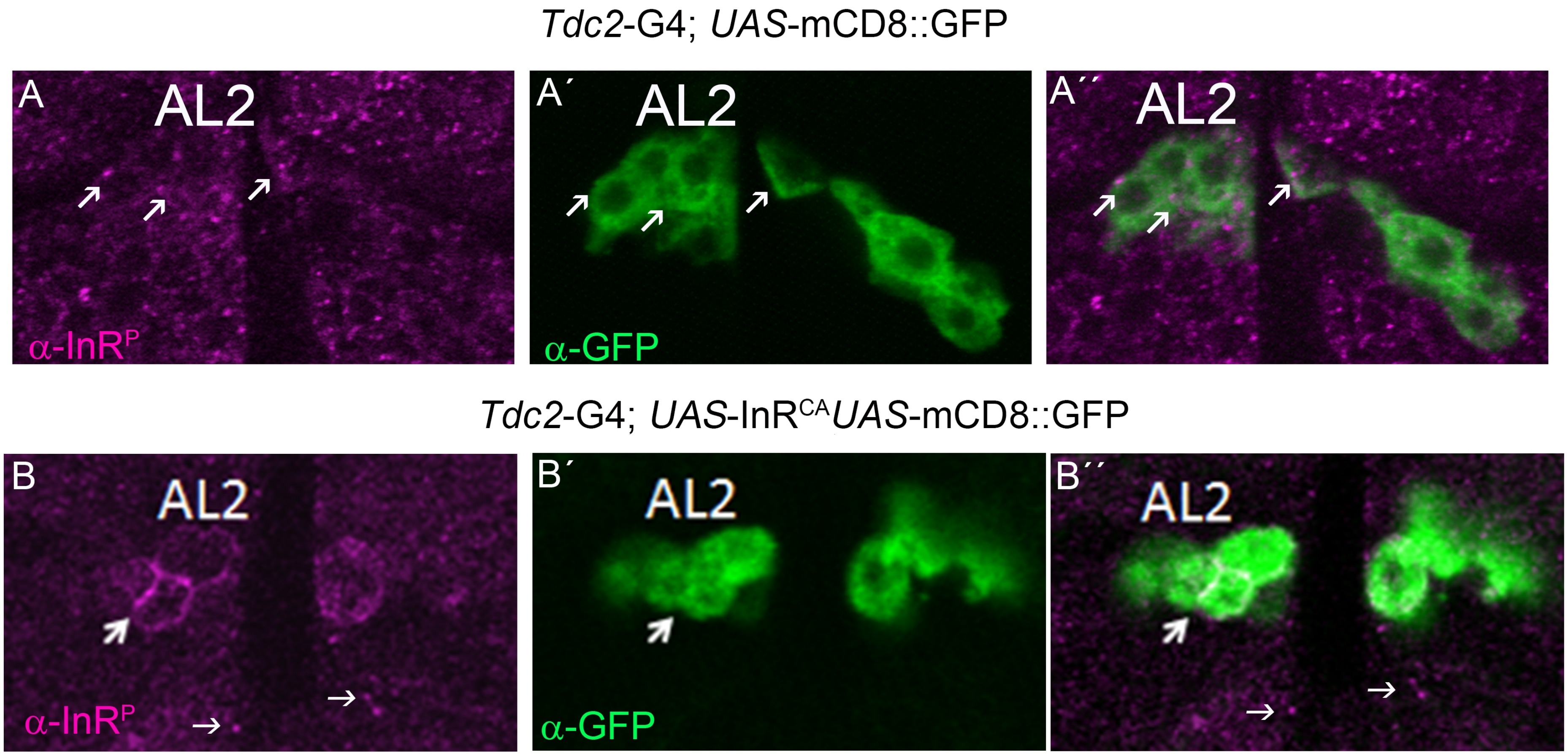
Tdc2-G4; UAS-mCD8::GFP
A
AL2
α-InR^P
A´
AL2
α-GFP
A´´
AL2
Tdc2-G4; UAS-InR^CA UAS-mCD8::GFP
B
AL2
α-InR^P
B´
AL2
α-GFP
B´´
AL2
